# Protein-based RBD-C-tag COVID-19 Vaccination Candidate Elicits Protection Activity against SARS-COV-2 Variant Infection

**DOI:** 10.1101/2021.08.17.456704

**Authors:** Jaganathan Subramani, Namir Shaabani, Dhwani Shetty, Haiwa Wu, Sunkuk Kwon, Wenzhao Li, Chanyu Yue, Christoph Lahtz, Adela Ramirez-Torres, Heyue Zhou, Yanliang Zhang, Robert Allen, Bill Farley, Mark Emalfarb, Ronen Tchelet, Soloheimo Markku, Vitikainen Marika, Marilyn Wiebe, Anne Huuskonen, Hannah Ben-artzi, Avi Avigdor, Henry Ji, Andreas Herrmann

**Author notes:** Contributed equally (Jaganathan Subramani, Namir Shaabani). Correspondence: Email Dr. rer. nat. Andreas Herrmann.

## Abstract

The identification of a vaccination candidate against COVID-19 providing protecting activity against emerging SARS-COV-2 variants remains challenging. Here, we report protection activity against a spectrum of SARS-COV-2 and variants by immunization with protein-based recombinant RBD-C-tag administered with aluminum-phosphate adjuvant intramuscularly. Immunization of C57BL/6 mice with RBD-C-tag resulted in the *in vivo* production of IgG antibodies recognizing the immune-critical spike protein of the SARS-COV-2 virus as well as the SARS-COV-2 variants alpha (“United Kingdom”), beta (“South Africa”), gamma (“Brazil/Japan”), and delta (“India”) as well as *wt*-spike protein. RBD-C-tag immunization led to a desired Th1 polarization of CD4 T cells producing IFNγ. Importantly, RBD-C-tag immunization educated IgG production delivers antibodies that exert neutralizing activity against the highly transmissible SARS-COV-2 virus strains “Washington”, “South Africa” (beta), and “India” (delta) as determined by conservative infection protection experiments *in vitro.* Hence, the protein-based recombinant RBD-C-tag is considered a promising vaccination candidate against COVID-19 and a broad range of emerging SARS-COV-2 virus variants.

## INTRODUCTION

Global efforts have been made to identify a vaccination candidate against infection with SARS-COV-2 virus causing COVID-19. However, a pluripotent antigen serving successful immunization against SARS-COV-2 and its variants emerging worldwide is imperative but remains challenging.

Vaccination strategies against SARS-COV-2 targeting the full-length viral spike glycoprotein range from employing attenuated virus and/or virus debris^1^ to DNA^2^ and RNA formats^3, 4^ encoding critical antigen to trigger a desired immune response. In addition, frequently challenging low antigen immunogenicity often requires contributing adjuvant to propagate a humoral response amplitude providing protective activity^5^. Moreover, particularly DNA and RNA vaccination formats involve nanoparticle technology to facilitate intracellular delivery^6^. Here, we employed recombinant RBD-C-tag protein to determine its immunization capacity against SARS-COV-2 and SARS-COV-2 variant infection.

The expression of the spike protein is an immunological hallmark of the SARS-COV-2 virus^7^. Emerging SARS-COV-2 variants are defined by distinct repertoires of point mutations in the viral spike-encoding gene^8, 9^ The mutation repertoires are unique for every virus variant such as the “United Kingdom” strain^10^, the “South African” strain^11^, the “Brazil/Japan” strain^12^, and the “India” strain^13^ and favor aggressive transmissibility to humans. The spike glycoprotein is largely composed of an extremely variable N-terminal S1 domain^14^ followed by a highly conserved S2 domain^15^ and a C-terminal transmembrane stretch of amino acids anchoring the spike protein onto the surface of viral membrane^7^. Pioneering crystallographic studies have demonstrated that the spike glycoprotein forms a trimer-like structure on the surface of viral membranes^16^. The receptor-binding domain (referred to as RBD) is integral part of the S1 domain of the spike protein^17^. RBD is thought to initiate SARS-COV-2 virus infection by interaction with host ACE2 receptors^17^ in humans while the S2 domain is emphasized to facilitate membrane fusion of virus and host cell membranes enabling viral cargo delivery into host cells^15^. Vaccination strategies targeting full-length spike glycoprotein are driven by the promise of *in vivo* educated IgG antibody populations binding to the spike protein and inhibit infection by preventing the critical spike-ACE2 interaction and subsequent cell infection^1,3, 4, 18^.

Here, we show that intramuscular administration into the *biceps femoris* of mice leads to accumulation of RBD-C-tag in the inguinal lymph node in conjunction with engagement and/or cellular internalization by immune cells of the antigen presenting compartment such as CD19^+^ B cells, CD11c^+^ dendritic cells, and CD11b^+^CD11c^-^ myeloid macrophages. Immunization of mice with RBD-C-tag in combination with aluminum adjuvant elicits the *in vivo* production of IgG antibody populations recognizing RBD as well as wt-S1 of the spike glycoprotein. Moreover, immunization with RBD-C-tag propagates IgG antibody populations recognizing a broad spectrum of spike glycoproteins including spike variants specifically identifying SARS-COV-2 virus alpha (“United Kingdom”), beta (“South Africa”), gamma (“Brazil/Japan”), and delta (“India”) as well as *wt*-spike protein (“Wuhan/Washington”). Interestingly, RBD-C-tag immunization educates a CD4^+^IFNγ^+^ Th1 T cell polarization, but hardly produces an IFNγ^+^ CD8 T cell population. However, triple immunization with RBD-C-tag 21 days apart in conjunction with aluminum phosphate adjuvant results in a broad and highly effective protection capacity by *in vivo* produced IgG antibodies, inhibiting infection by SARS-COV-2 virus strain beta (“South Africa”), gamma (“Brazil/Japan”), and delta (“India”) as well as the SARS-COV-2 virus prototype strain (“Wuhan/Washington”). Hence, the SARS-COV-2 vaccination candidate rRBD-C-tag represents an operative immunization approach against COVID-19 exerting multifaceted protection activity.

## RESULTS

### RBD-C-tag homes to lymph nodes *in vivo* and undergoes cellular internalization by immune cells

The biodistribution of the vaccination candidate RBD-C-tag was monitored upon administration. Therefore, RBD-C-tag was fluorescently labeled with a fluorophore that emits light in the near infra-red spectrum to facilitate non-invasive *in vivo* imaging. Fluorescently labeled RBD-C-tag^IRDye800^ was injected intramuscularly into the *biceps femoris* of C57BL/6 mice and whole-body imaging was carried out immediately after injection. *In vivo* delivered RBD-C-tag was robustly detectable in the inguinal lymph node after 1 hour of injection in a dose-dependent manner (Fig. 1A). Notably, RBD-C-tag did not undergo immediate lymphatic clearance but accumulated in the inguinal lymph node for up to 4 hours upon administration indicative of cellular engagement and/or internalization. *In vitro,* RBD-C-tag was shown to undergo cellular internalization by splenic CD11b^+^ and CD11c^+^ myeloid antigen-presenting cells, CD3^+^ T cells and CD19^+^ B cells favoring the orchestration of a crucial RBD-antigen-dependent immune response (Fig. 1B).

**Figure 1:**
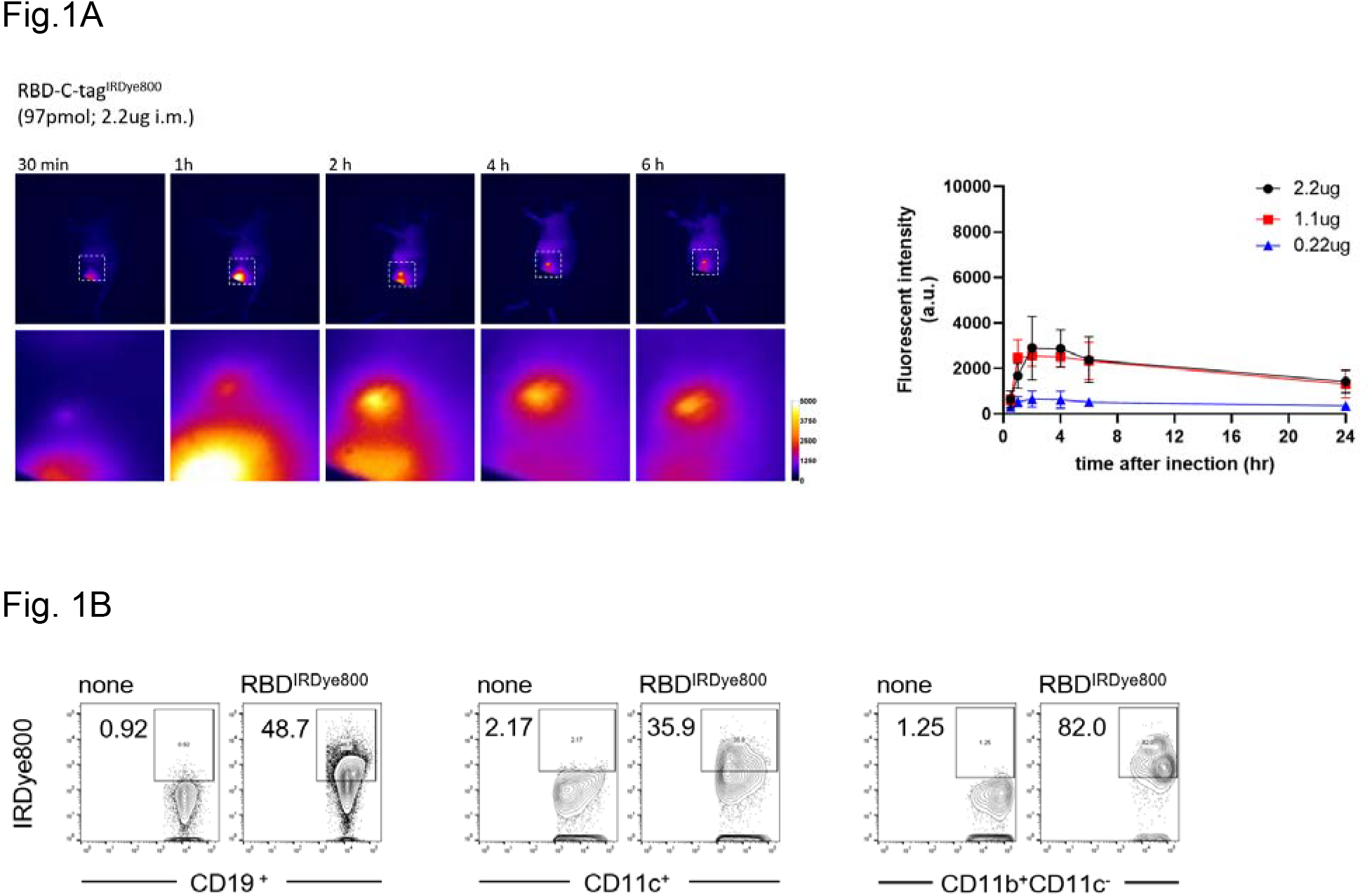
RBD-C-tag homes to the draining lymph node and undergoes uptake by antigen-presenting cells. **A**, Fluorescently labeled RBD-C-tag was administered intramuscularly and live imaging was carried out to determine and quantify homing and lymphoid accumulation in a dose-dependent manner. SD shown. **B**, Cellular uptake of fluorescently labeled RBD-C-tag was assessed using freshly isolated splenic immune cell subsets CD19 B cells, CD11c dendritic cells and CD11b^+^CD11c^-^ myeloid macrophages.

### RBD-C-tag immunization induces anti-spike IgG production *in vivo*

The receptor-binding domain (RBD) at the membrane-distal tip of the viral spike protein is critical for the engagement of the SARS-COV-2 virus to engage with host-recipient receptor ACE2. Hence, blocking the RBD/ACE2 interaction by RBD-specific antibodies represents an enticing concept to prevent virus infection. Here, RBD-C-tag was employed as an antigen to raise an antigen-specific immune response against viral RBD. Notably, RBD formats are of low immunogenicity (*data not shown*) and require adjuvant to induce a desired immune response. This prompted us to include, evaluate and compare the immune response to RBD in the presence of aluminum hydroxide adjuvant or aluminum phosphate adjuvant. Moreover, a second “booster” immunization (3 immunizations total) considered to elevate anti-RBD IgG production in vivo was included and compared to two immunizing administrations at high (100 μg) or low (50 μg) doses (Fig. 2A, B).

**Figure 2:**
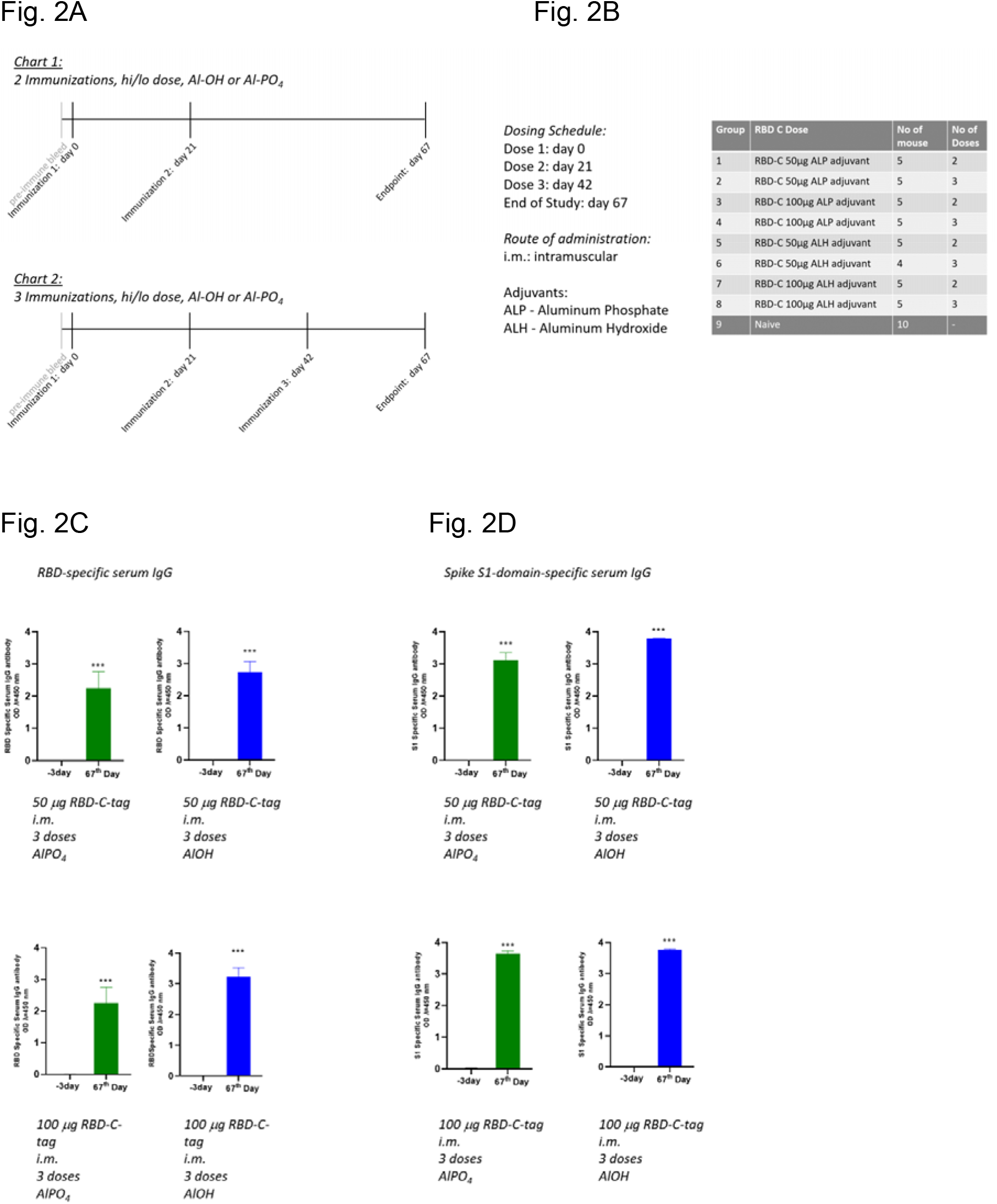
Immunization with RBD-C-tag propagates production of RBD- and spike-S1-specific antibodies *in vivo*. **A**, Immunization regimen over the course of 67 days with hi/lo dose immunization, 2 or 3 immunizations and comparing the adjuvants aluminum hydroxide and aluminum phosphate (**B**) unfolds in 8 mice cohorts. **C**, RBD-specific serum IgG and (**D**) spike S1-subdomain-specific serum IgG was assessed by ELISA. SD shown. T-test: ***) *P*<0.001.

Immunization with RBD-C-tag in the presence of aluminum adjuvants induced the production of RBD-specific IgG antibodies *in vivo* (Fig. 2C). Interestingly, neither the single dose level nor the adjuvants employed facilitated a distinct IgG antibody production *in vivo* but favored a similar IgG antibody production. Moreover, *in vivo* produced IgG antibodies recognized RBD in the context of the S1 subdomain of the wild-type spike protein independent of administration regimen (Fig. 2D), which is critical to predict an inhibitory and/or neutralizing capacity of IgG antibodies produced upon immunization.

### SARS-COV-2 variant recognition upon RBD-C-tag immunization

Although emerging SARS-COV-2 variants have been shown to undergo extensive mutagenesis in the spike-protein encoding gene, it is to be anticipated that immunization with protein-based formats will inevitably result in the production of a polyclonal antibody population *in vivo* with the beneficiary potential of a wide range of inhibition modalities, blocking the RBD/ACE2 interaction at multiple sites. Immunization with RBD-C-tag in the presence of adjuvants resulted in the production of an antibody population that recognizes a broad range of SARS-COV-2 spike protein variants (Fig. 3A). As assessed by ELISA, in vivo produced IgG antibodies recognize the SARS-COV-2 alpha variant (“United Kingdom”), the beta variant (“South Africa”), the gamma variant (“Brazil/Japan”), and the delta variant (“India”). Again, the production of anti-spike IgG antibodies was detected similar and apparently independent on the dose administered or the adjuvant present. Interestingly, high dilution ranges of anti-pan-spike seropositive mouse blood serum are indicative of a high immunogenic potential of RBD-C-tag (Fig. 3B). Since the spike protein has been described to form trimers on the surface of the viral membrane^16^, the binding of murine IgG antibodies to spike protein variants was assessed in a cell-based assay to allow a viral-like spike-trimer formation and mimic a more physiologic condition. Hence, spike and/or mutated spike proteins were overexpressed in human Hek293 cells. Murine blood serum seropositive for IgG antibodies produced *in vivo* upon RBD-C-tag immunization was diluted and incubated with spike protein expressing Hek293 cells. Murine IgG antibodies produced *in vivo* upon RBD-C-tag immunization could be demonstrated to engage with Hek293 cells expressing SARS-COV-2 wt-spike, the alpha spike variant, the beta spike variant, and the gamma spike variant as determined by flow cytometry (Fig. 3C). However, IgG antibodies produced *in vivo* did not engage with parental Hek293 cells which served as an experimental control. Thus, immunization using RBD-C-tag and aluminum adjuvants induces the production of a serum IgG population that recognizes a wide range of SARS-COV-2 spike proteins and represents an omnipotent vaccination candidate.

**Figure 3:**
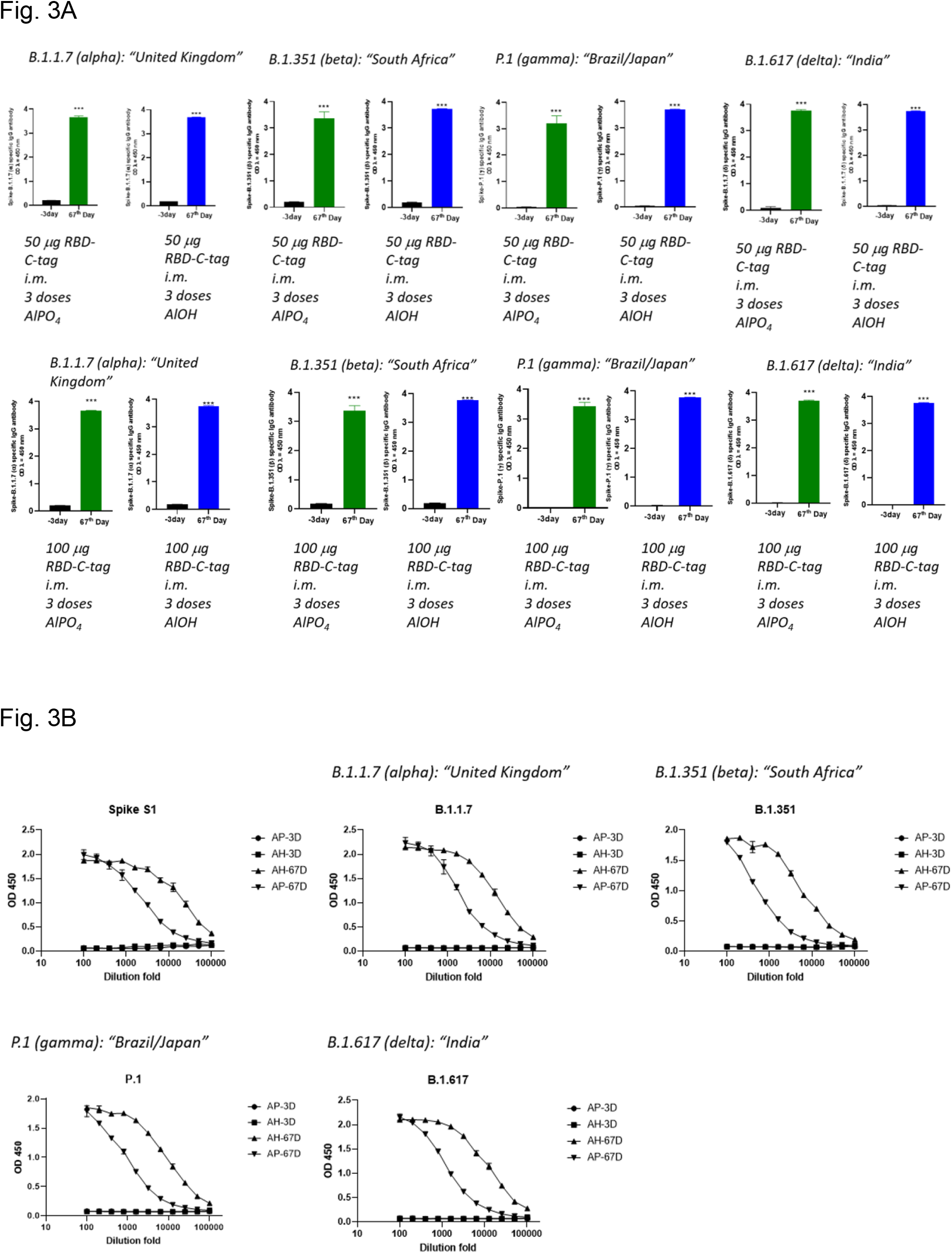
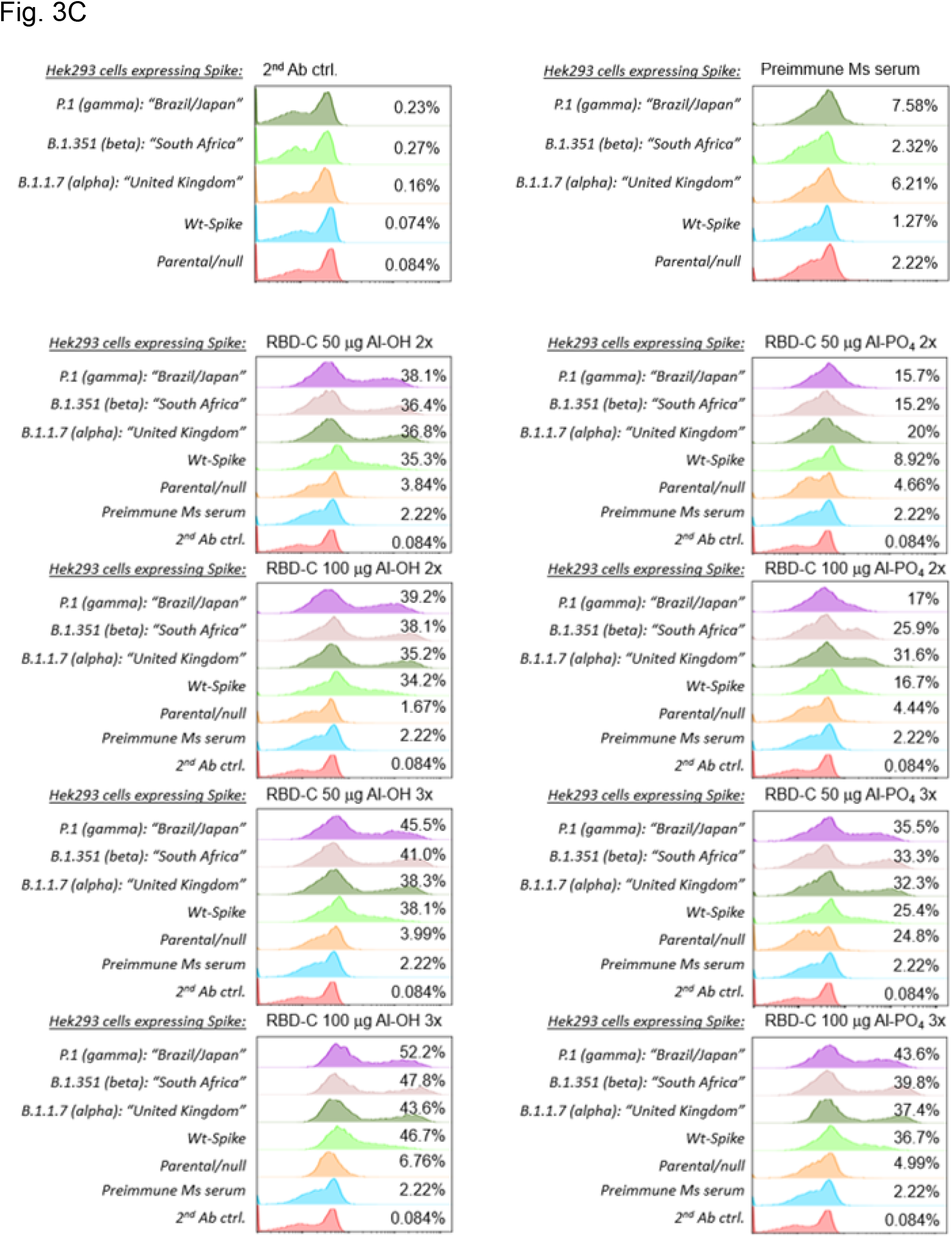
RBD-C-tag immunization facilitates *in vivo* production of serum IgG antibodies recognizing the broad spectrum of SARS-COV-2 variants. **A**, Pre-immune mouse serum and seropositive blood serum isolated after immunizations (day 67) were compared for serum IgG specificity recognizing spike proteins representing SARS-COV-2 variants by ELISA, SD shown. T-test: ***) *P*<0.001. **B**, *In vivo* produced serum IgG was tested for antigen recognition potency by serial dilution; pre-immune serum was included as a control in ELISA-based assessment. **C**, SARS-COV-2 spike variant recognition by serum IgG was assessed in a cell-based flow cytometric assay using Hek293 cells expressing the viral spike protein variants, potentially allowing spike protein trimer formation on the surface of cells in a viral-like fashion. Parental Hek293 cells not expressing spike protein were included as a negative control.

### RBD-C-tag immunization favors an IFNγ producing Th1 T cell polarization

To assess immune cell responses upon RBD-C-tag immunization in the presence of adjuvants, single cell populations were prepared from spleens of immunized mice at the experimental end-point day 67. Notably, immunization with RBD-C-tag favored the maturation of IFNγ producing CD3^+^CD4^+^ T cells indicative of a Th1 polarization (Fig. 4A). However, effector IFNγ producing CD8^+^ T cells (Fig. 4B) were hardly detectable, and peripheral IL-10 producing CD3^+^CD4^+^ Treg cell/IL-17 producing CD3^+^CD4^+^ Th17 cell plasticity was not regulated (Fig.4C) upon RBD-C-tag immunization. Hence, RBD-C-tag immunization must be considered to educate a rather mild T cell response, yet in favor of an anti-viral immune response by IFNγ production.

**Figure 4:**
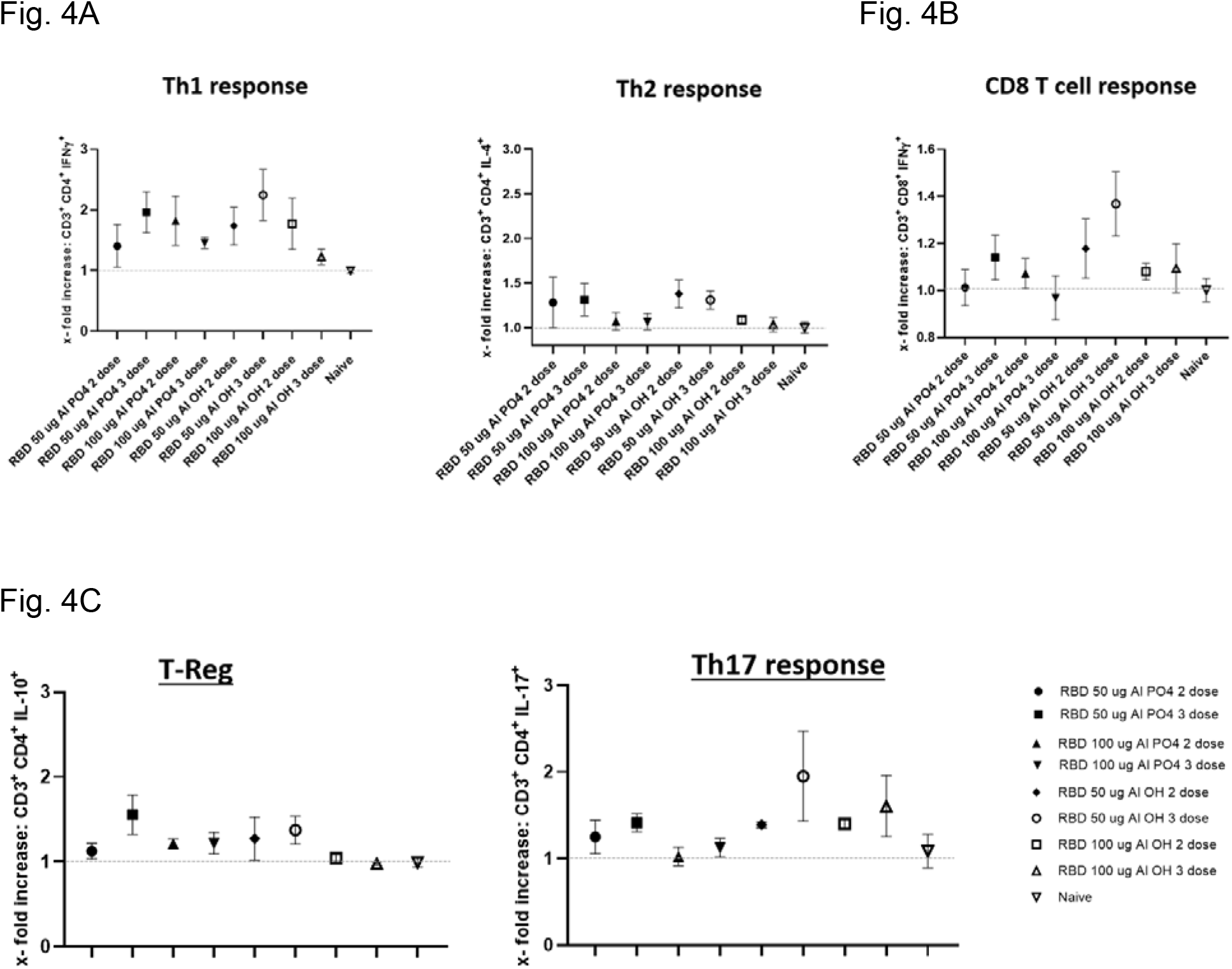
Immunization with RBD-C-tag propagates a CD4^+^ Th1 T cell polarization *in vivo*. Splenic single sell populations were isolated after immunizations and (**A**) CD3^+^CD4^+^IFNγ^+^ Th1 cell maturation, CD3^+^CD4^+^IL-4^+^ Th2 cells maturation, as well as (**B**) CD3^+^CD8^+^IFNγ^+^ effector CD8 T cells were assessed by flow cytometry. (**C**) In addition, CD3^+^CD4^+^IL-10^+^ regulatory T cells and CD3^+^CD4^+^IL-17^+^ Th17 T cell plasticity was assessed. Splenic cell populations isolated from naïve mice were included as a reference control.

### RBD-C-tag immunization elicits protection against SARS-COV-2 infection and SARS-COV-2 variant infection *in vitro*

The amplitude of protection against SARS-COV-2 infection and SARS-COV-2 variant infection was assessed in a conservative PRNT assay *in vitro*, where healthy VeroE6 cells were directly exposed to live virus in the absence or presence of diluted mouse blood serum seropositive for anti-spike IgG antibodies. PRNT detected plaque formation is indicative of cell infection by virus while reduced plaque formation informs the neutralizing capacity of serum IgG antibodies produced upon RBD-C-tag immunization. Notably, employing the adjuvant aluminum phosphate as well as an additional second “booster” immunization (3 immunizations total) at elevated dosing are decisive to induce the production of a potent IgG antibody population sufficing protection against SARS-COV-2 (Fig. 5A). As shown by PRNT assay, 3 doses of 100 μg RBD-C-tag 21 days apart, intramuscular administration, and the presence of aluminum phosphate raise neutralizing antibodies, protecting VeroE6 cells from infection by either the “Washington/Wuhan” (strain of origin) SARS-COV-2 strain, the “South Africa” (beta) strain, and the “India” (delta) strain in a conservative experimental setting. Protection against the “Brazil/Japan” (gamma) SARS-COV-2 strain reached approximately 60% efficacy as assessed using a 1:40 dilution of blood sera isolated from immunized mice. Although the adjuvant aluminum hydroxide contributed to induce a promising anti-spike serum IgG population, its protection amplitude is rather low when compared to immunization using aluminum phosphate. In addition, further serial dilution of blood sera informed the potency of IgG antibodies propagated by immunization with RBD-C-tag indicating robust protection activity against the “Washington/Wuhan” (strain of origin) SARS-COV-2 strain, the “South Africa” (beta) strain, and to a lesser extend against the “India” (delta) strain (Fig. 5B). Thus, RBD-C-tag immunization contributed by aluminum phosphate adjuvant has shown a potent protection amplitude against SARS-COV-2 infection and SARS-COV-2 variant infection.

**Figure 5:**
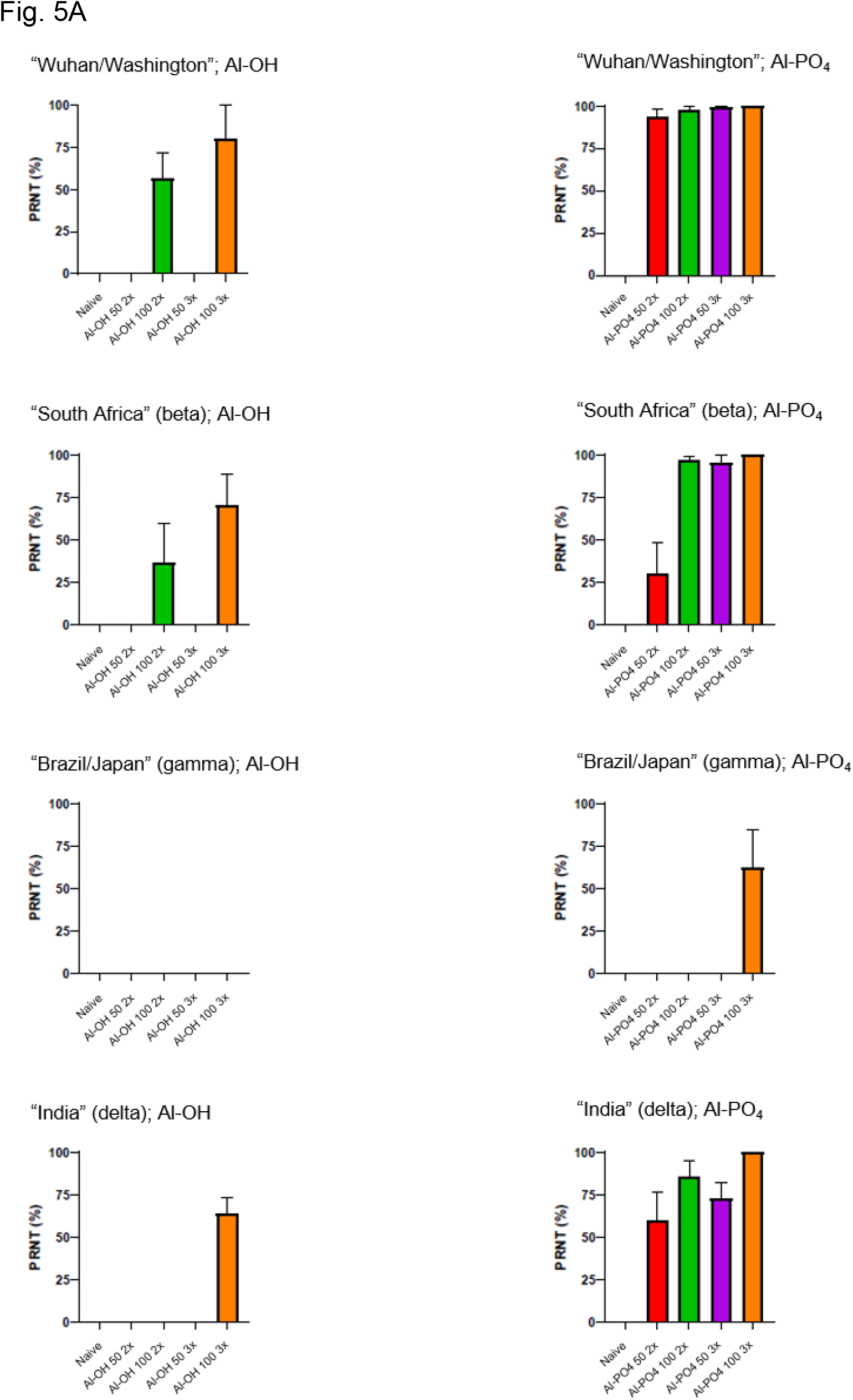
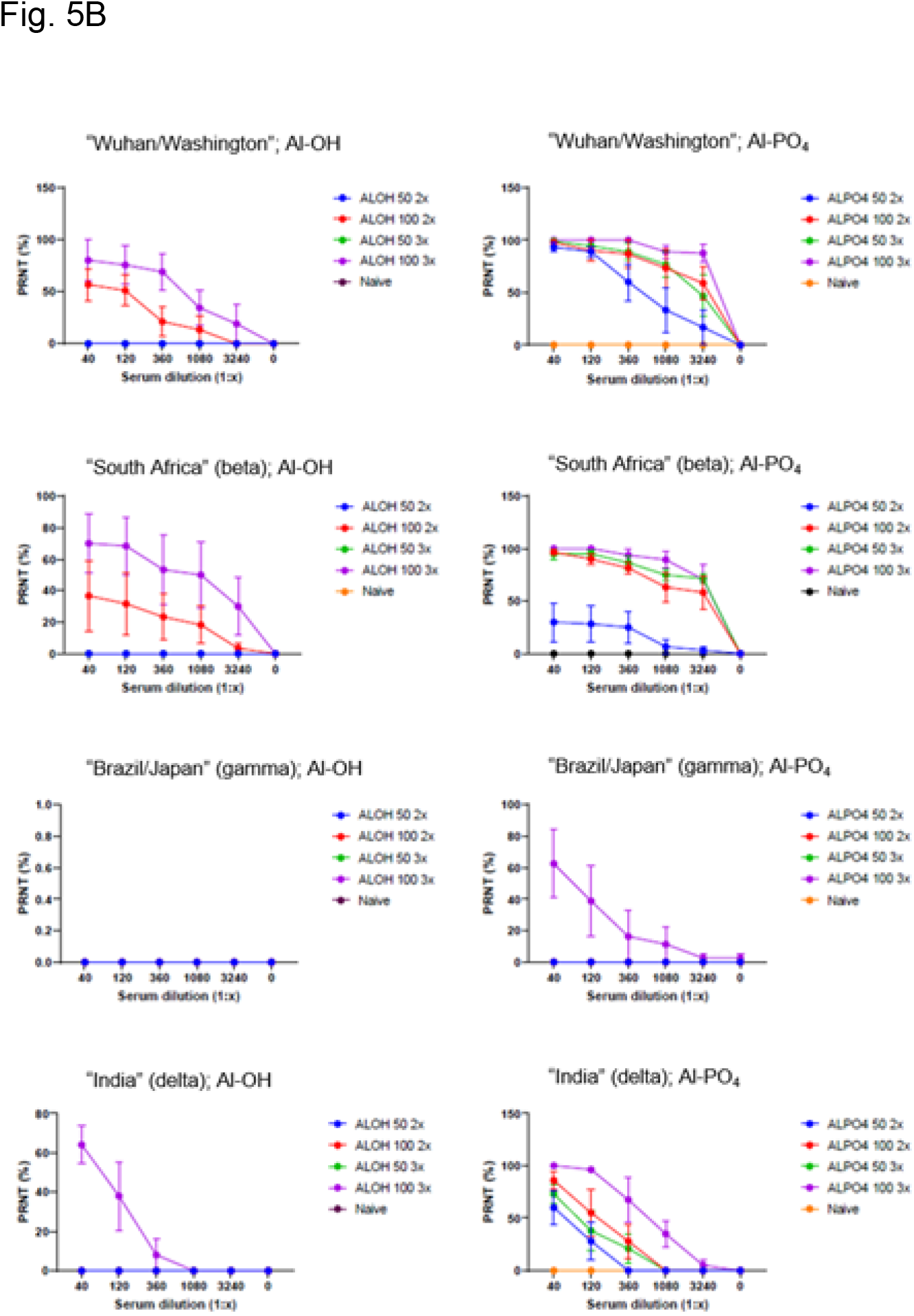
RBD-C-tag immunization achieves a potent protection amplitude against SARS-COV-2 and SARS-COV-2 variant infection. As determined by a conservati**ve** live virus challenge assessment *in vitro*, PRNT assays were carried out (**A**) at a 1:40 dilution of blood sera isolated from mice and/or (**B**) at serial dilutions of blood sera isolated from mice was performed to estimate the potency of serum IgG produced **in** vivo upon RBD immunization. SD shown.

## DISCUSSION

Of the three virus outbreaks in humans caused by *betacoronaviridae* since 2002, the SARS-COV-2 driven COVID-19 pandemic owes the highest toll and is associated with 177 million infection cases worldwide and 3.8 million deaths as of June 2021^19^ In 2002/2003, the SARS-CoV outbreak was reported with approximately 9,000 cases associated with 800 deaths. In 2012, the Middle-East Respiratory Syndrome caused by MERS-CoV counted 3,000 cases in total^20^.

Vaccination campaigns against SARS-COV-2 causing COVID-19 employing mRNA formats and/or adenoviral backbone entailing formats showed promising efficacy but must be interrogated for persistence as well as for protection capacity while new SARS-COV-2 virus variants are emerging.

Here, we report a protein-based COVID-19 vaccination candidate RBD-C-tag, that induces the production of a multivalent IgG antibody population when administered three times in combination with the adjuvant aluminum phosphate. Interestingly, the aluminum phosphate adjuvant as well as adding a second “booster” immunization considerably improved the vaccination efficacy as indicated by the increased protection capacity determined by PRNT assay. However, further testing of adjuvants such as saponin-based adjuvant, TLR agonists/CpG-containing oligonucleotides and/or additional aluminum salts might be considered supportive in the effort to decrease the immunization single-dose, potentially reduce the frequency of dosing, and lessening the vaccination burden of patients.

An omnipotent immunity raised by a safe multivalent COVID-19 vaccination candidate is desperately needed to avoid perpetual immunization campaigns directed by newly emerging SARS-COV-2 variants. Currently distributed and administered COVID-19 vaccinations have been shown to elicit limited protection activity^21,22^. Since the common scientific opinion largely agrees on the targeting mechanism, the immunizing antigen is defined on the molecular level^3, 4, 18^. However, the therapeutic benefit of a multivalent SARS-COV-2 immunization depends on the design, production/manufacturing, and administration route of the vaccination candidate. Hence, additional supporting studies are required to identify an optimized administration route, considering that intramuscular administration is just one option amongst a variety of routing technologies including but not restricted to lymphatic delivery^23, 24^. Thus, the optimized administration of a multivalent SARS-COV-2 vaccination candidate potentially could contribute to adjust and/or augment previously limited immunization capacity and broaden the range of protection.

Various viruses belonging to the family of *coronoviridae* are known to typically cause rather mild self-limited infections located in the upper respiratory tract, common colds, infectious bronchitis associated with worldwide virus prevalence and circulating in human populations since a long time, but also severe acute respiratory syndrome (SARS)^25, 26^. Going forward, similar to the immunizing control of influenza virus contraction, a routine seasonal SARS-COV-2 vaccination might be considered to overcome and control SARS-COV-2 infection. Hence, continuous monitoring will inevitably enhance our understanding of newly evolved virus variants and eventually guide to control of the current SARS-COV-2 pandemic.

## MATERIALS AND METHODS

### Mice and cell culture

For intramuscular antigen challenge, C57BL/6 female mice 5-6 weeks old were injected with either 100 μg RBD-C-tag (300 μg adjuvant) or 50 μg RBD-C-tag (150 μg adjuvant) intramuscularly. Antigen challenge at days according to 2 immunizations or 3 immunizations was administered 21 days apart into the *biceps femoris* muscles of mice. Peripheral blood was collected from anaesthetized mice once/week via retro-orbital route. Simian VeroE6 cells were obtained from ATCC (CRL-1586) and subcultured in DMEM supplemented with 10% FBS. Human parental HEK293 cells (ATCC, CRL-1573) were cultured in DMEM supplemented with 10% FBS and antibiotics/antimycotics (Gibco).

### RBD-C-tag and adjuvants

Recombinant RBD-C-tag was produced by Dyadic International (Jupiter, FL) using an established expression system based on the fungus *Myceliophthora thermophila*. Aluminum adjuvants aluminum phosphate (InvivoGen; vac-phos-250) and aluminum hydroxide (InvivoGen; vac-alu-250) were used in the ration in the ratio of 1:1 v/v.

### Generation of Hek293 cells stably expressing SARS-COV-2 spike-protein variants

Human parental HEK293 cells were cultured in DMEM supplemented with 10% FBS. Plasmids encoding the SARS-COV-2 spike glycoprotein or its mutated variants were cloned into a G418 resistance gene containing backbone and transfected into HEK293 cells using Lipofectamine 2000 (Thermofisher) according to manufacturer’s instructions. 24-48 h after transfection, transgenic Hek293 cells were selected with 0.5 mg/ml G418 for 2-3 weeks before colonies were picked and expanded.

### Fluorescent labeling of RBD-C-tag

RBD-C-tag was conjugated to IRDye^®^ 800RS NHS Ester (Li-cor) using amine-reactive crosslinker chemistry. Briefly, RBD-C-tag antigen (2.5-3.0 mg/ml) was reacted with 5 equivalents of IRDye^®^ 800RS NHS Ester in 1× DPBS pH 7.4 containing 5% anhydrous DMSO for 3 hours with gentle rotation at room temperature. Subsequently, the reaction mixture was subjected to PD-Minitrap G-25 column (GE Healthcare) to remove unreacted dyes according to the manufacturer’s instructions. Upon purification, the conjugate underwent buffer exchange three times into 1× DPBS pH 7.4 using a 4-ml Amicon Ultra centrifugal filter (30 kDa MWCO, Millipore). The conjugate was characterized using SDS-PAGE, SEC HPLC, and BCA Assay.

### Longitudinal near infrared in vivo imaging of lymphatics

IRDye800-RBD-C-tag was injected intramuscularly into the *biceps femoris* of C57BL/6 mice. Near-infrared fluorescence imaging kinetics at 100 ms exposure time was performed upon administration. The injection site was concealed to prevent photonic overexposure. Fluorescent RBD-C-tag homing to/clearing from the inguinal lymph node was quantified by applying a region of interest (ROI) to determine fluorescent intensity kinetics.

### Detection of RBD and Spike binding serum IgG antibodies by ELISA

A direct binding ELISA format was used to detect the anti-SARS-COV-2 RBD or Spike IgG antibody in mouse serum samples. The plate was coated with SARS-COV-2 (2019-nCoV) Spike Protein (S1 Subunit; Gene script, Z03501-1) or RBD-C tag (Dyadic International, Jupiter, FL) or different variants of spike protein B.1.1.7 (Alpha; Cube Biotech # 28718), B.1.351 (Beta; Cube Biotech # 28721), P.1 (Gamma; Cube Biotech #28723), and B.1.617.2 (Delta; Cube Biotech #28741) at 5 μg/ml at 4°C overnight. The next day, the plate was washed 3 times with 1× KPL buffer (Sera Care, # 5150-0008) and blocked with Superblock (Scy Tek Labs #AAA500) at room temperature (RT) for 1 h. Mouse serum samples were diluted with Casein Block Buffer at the designated dilution factor. The blocked plate was washed once and incubated with the samples at RT for 1.5 h shaking at 150 rpm. The plate was washed 3 times before adding 50 μl of 1:1,000 diluted Goat anti-Mouse IgG H+L-HRP antibody (Invitrogen, # 31430) to the plate and incubated for 1 h at RT, 150 rpm. The plate was washed 3 times before adding 50 μl of KPL TMB substrate (Sera care, # 5120-0075) to each well. The plate was incubated at RT for 10-15 min. At the end of incubation, 50 μl of stop solution (Invitrogen, # REF SS04) was added to each well to stop TMB development, and optical density OD was assessed at λ=450 nm immediately.

### Intracellular Staining and Flow Cytometry

To prepare single-cell suspensions for flow cytometry, spleens from naïve or immunized mice were collected and homogenized in PBS^27^. Digests were filtered through 70 μm cell strainers, centrifuged at 1,500 rpm for 5 min. After red blood cell lysis (Biolegend; # 420301), single-cell suspensions were filtered, washed, and resuspended in RPMI with 10% FBS medium and were stimulated for 6 h with a cell activation cocktail containing PMA and ionomycin (Biolegend, # 423302) in the presence of protein transport inhibitor, Brefeldin A (Biolegend, #420601). At the end of the stimulation, the cells were washed, fixed with 2% PFA, permeabilized with ice-cold methanol, blocked with CD16/CD32, and stained for 30 min on ice with CD3 FITC (Biolegend, # 100306), CD4 BV421 (Biolegend, # 100438), CD8c BV650 (Biolegend, #100742), IFNγ PE (Biolegend, # 505808), IL-4 APC (Biolegend, #504106), IL-17 PECy7 (Biolegend, # 506922), donkey anti-mouse IgG conjugated to AlexaFluor 647 (Invitrogen #A-31571) or IL-10 APC antibodies ( eBioscience, # 17-7101-81). Cells were washed twice before analysis on the BD FACS*Celesta* flow cytometer (Becton Dickinson).

### Plaque Reduction Neutralization Test (PRNT)

Simian VeroE6 cells were plated at 18×10^3^ cells/well in a flat bottom 96-well plate in a volume of 200 μl/well. After 24 h, a serial dilution of seropositive blood serum is prepared in a 100 μl/well at twice the final concentration desired and live virus was added at 1,000 PFU/100μl of SARS-COV-2 or SARS-COV-2 variants (BEI Resources) and subsequently incubated for 1 h at 37°C in a total volume of 200 μl/well. Cell culture media was removed from cells and sera/virus premix was added to VeroE6 cells at 100 μl/well and incubated for 1 h at 37°C. After incubation, 100 μl of “overlay” (1:1 of 2% methylcellulose (Sigma) and culture media) is added to each well and incubation commenced for 3 d at 37°C. Plaque staining using Crystal Violet (Sigma) was performed upon 30 min of fixing the cells with 4% paraformaldehyde (Sigma) diluted in PBS. Plaques were assessed using a light microscope (Keyence).

### Study approval

Mouse care and experimental procedures with mice were performed under pathogen-free conditions in accordance with established institutional guidance and approved Animal Care and Use Protocols (ACUP) from the Research Animal Care Committee at Sorrento Therapeutics Inc.

## Notes

### Competing Interest Statement

The authors have declared no competing interest.

